# Using DeepLabCut for 3D markerless pose estimation across species and behaviors

**DOI:** 10.1101/476531

**Authors:** Tanmay Nath, Alexander Mathis, An Chi Chen, Amir Patel, Matthias Bethge, Mackenzie Weygandt Mathis

## Abstract

Noninvasive behavioral tracking of animals during experiments is crucial to many scientific pursuits. Extracting the poses of animals without using markers is often essential for measuring behavioral effects in biomechanics, genetics, ethology & neuroscience. Yet, extracting detailed poses without markers in dynamically changing backgrounds has been challenging. We recently introduced an open source toolbox called DeepLabCut that builds on a state-of-the-art human pose estimation algorithm to allow a user to train a deep neural network using limited training data to precisely track user-defined features that matches human labeling accuracy. Here, with this paper we provide an updated toolbox that is self contained within a Python package that includes new features such as graphical user interfaces and active-learning based network refinement. Lastly, we provide a step-by-step guide for using DeepLabCut.

## Introduction

Advances in computer vision and deep learning are transforming research. Applications like autonomous car driving, estimating the poses of humans, controlling robotic arms, or predicting the sequence specificity of DNA- and RNA-binding proteins are rapidly developing [1–4]. However, many of the underlying algorithms are “data hungry” as they require thousands of labeled examples, which potentially prohibits these approaches to be useful for small scale operations, such as in single laboratory experiments [5]. Yet transfer learning, the ability to take a network, which was trained on a task with a large supervised data set, and utilize it for another task with a small supervised data set, is beginning to allow users to broadly apply deep learning methods [6–10]. We recently demonstrated that due to transfer learning, a state-of-the-art human pose estimation algorithm called DeeperCut [10, 11] could be tailored to use in the laboratory with relatively small amounts of annotated data [12]. Our toolbox, DeepLabCut [12], provides tools to create annotated training sets, train robust feature detectors, and utilize them to analyze novel behavioral videos. Current applications range across the common model systems like mice, zebrafish and flies as well as rarer ones like babies and cheetahs (Figure 1)

**FIG. 1.**
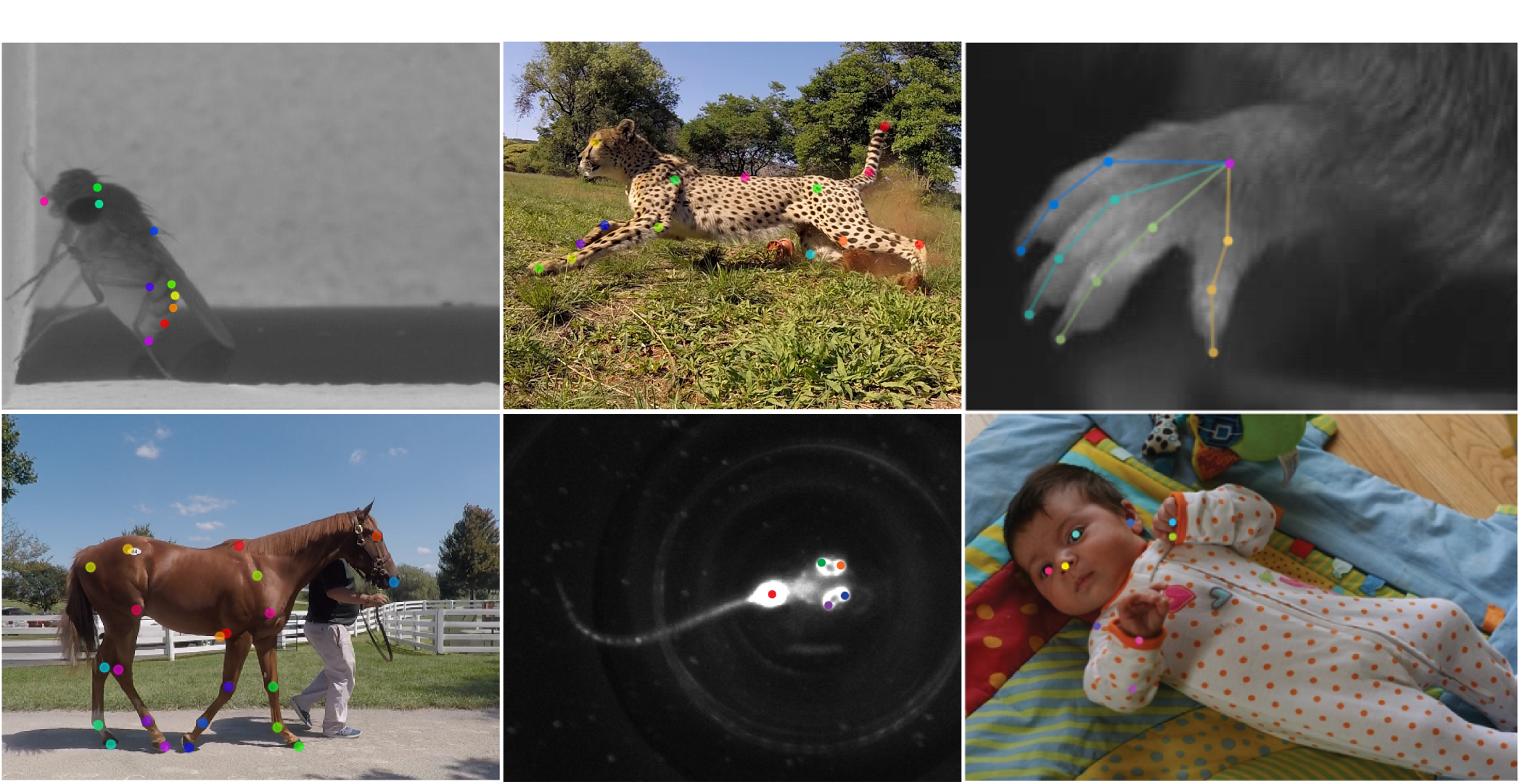
Pose estimation with DeepLabCut. Six examples with automatically applied labels from DeepLabCut. The colored points represent features the user wished to measure. Clockwise from the top left: a fruit fly, a cheetah, a mouse hand, a baby, a zebrafish, and a horse. The mouse hand image is adapted from [12], and the baby image is adapted from [26].

Here, we provide a comprehensive protocol and expansion of DeepLabCut that allows researchers to estimate the pose of the subject, efficiently enabling them to quantify the behavior. The major motivation for developing the DeepLabCut toolbox was to provide a reliable and efficient tool for high-throughput video analysis, where powerful feature detectors of user-defined body parts need to be learned for a specific situation. The toolbox is aimed to solve the problem of detecting body parts in dynamic visual environments where varying background, reflective walls or motion blur hinder the performance of common techniques, such as thresholding or regression based on visual features [13–21]. The uses of the toolbox are broad, however certain paradigms will most benefit from this approach. Specifically, DeepLabCut is best suited for behaviors that can be consistently captured by one or multiple cameras with minimal occlusions.

The DeepLabCut Python toolbox is versatile, and easy to use without extensive programming skills. With only a small set of training images a user can train a network to within human level labeling accuracy, thus expanding its application to not only behavior analysis in the laboratory, but to potentially also in sports, gait analysis, medicine and rehabilitation studies [13, 22–26].

## Overview of using DeepLabCut

DeepLabCut is organized according to the following workflow (Fig 2). The user starts by creating a new project based on a project and username as well as some (initial) videos, which are required to create the training dataset (Step A-B). Additional videos can also be added after the creation of the project, which will be explained in greater detail below. Next, DeepLabCut extracts frames, which reflect the diversity of the behavior with respect to postures, animal identities, etc. (Step C). Then the user can label the points of interest in the extracted frames (Step D). These annotated frames can be visually checked for accuracy, and corrected if necessary (Step E). Eventually, a training dataset is created by merging all the extracted labeled frames and splitting them into subsets of test and train frames (Step F). Then, a pre-trained network (ResNet) is trained end-to-end to adapt its weights in order to predict the desired features (i.e. labels supplied by the user; Step G). The performance of the trained network can then be evaluated on the training and test frames (Step H). The trained network can be used to analyze videos yielding extracted pose files (Step I). In case the trained network does not generalize well to unseen data in the evaluation and analysis step (Step H), then additional frames with poor results can be extracted (Step J) and the predicted labels can be manually shifted to their ideal location (Step K). This refinement step, if needed, creates an additional set of annotated images that can then be merged with the original training dataset to create a new training dataset (Step F). This larger training set can then be used to re-train the feature detectors for better results (Step G). This active learning loop can be done iteratively to robustly and accurately analyze videos with potentially large variability -i.e. experiments that include many individuals, and run over long time periods. Furthermore, the user can add additional body parts/labels at later stages during a project (Step D) as well as correct user-defined labels (Step K). Lastly, we provide Jupyter Notebooks (Section M), a summary of the commands (Section N), and a troubleshooting section (Section O).

**FIG. 2.**
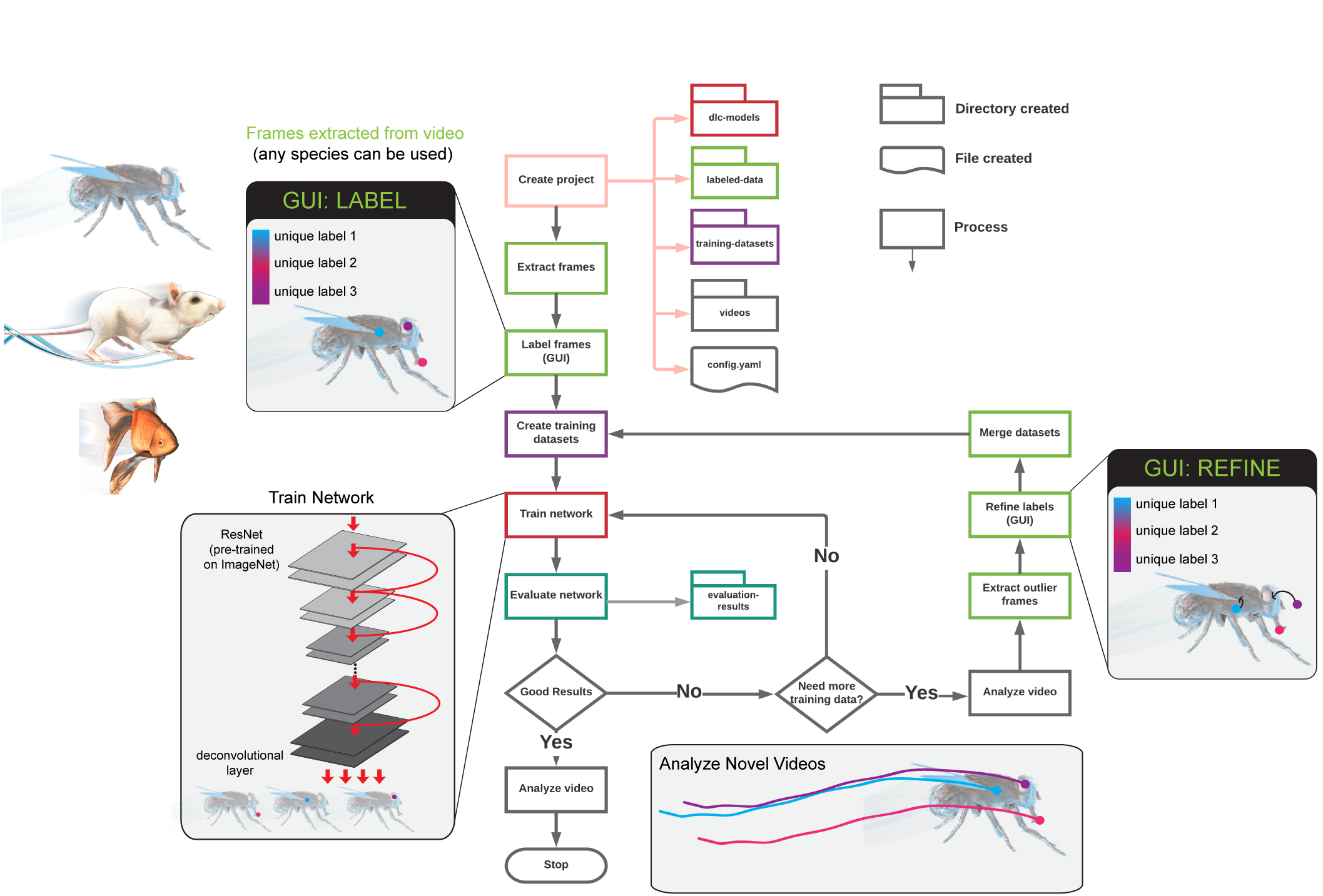
DeepLabCut work-flow. The diagram delineates the work-flow as well as the directory and file structures. The process of the work-flow is color-coded to represent the location of their output. The main steps are opening a python session, importing deeplabcut, creating a project, selecting frames, labeling frames, then training a network. Once trained, this network can be used to apply labels on new videos, or the network can be refined if needed. The process is enabled by interactive GUIs at several key steps, and all lines can be run from a simple terminal interface.

## Applications

DeepLabCut has been used to extract user-defined, key body part locations from animals (including humans) in various experimental settings. Current applications range from tracking mice in open-field behaviors, to more complicated hand articulations in mice, whole-body movements in flies in a 3D environment (all shown in [12]), as well as pose-estimation in human babies [26], 3D-locomotion studies in rodents [27], multi-bodypart including eye tracking during perceptual decision making [28], horses and cheetahs (Figure 1). Other (currently unpublished) use cases can be found at www.mousemotorlab.org/deeplabcut. Lastly, 3D pose estimation for a challenging cheetah hunting data set will be presented within this manuscript.

## Comparison with other methods

Pose-estimation is a challenging, yet classical computer vision problem [29], whose human pose estimation benchmarks have recently been shattered by deep learning algorithms [2, 10, 11, 30–33]. There are several main considerations when deciding on what deep learning algorithms to use, namely, the amount of labeled input data required, the speed, and the accuracy. DeepLabCut was developed to require little labeled data, to allow for real-time processing (i.e. as fast or faster than camera acquisition), and be as accurate as human annotators.

DeepLabCut utilizes parts of an algorithm called DeeperCut, which is one of the best algorithms for several human pose estimation benchmarks in recent years [10]. Specifically, DeepLabCut uses deep, residual networks with either 50 or 101 layers (ResNets) [34] and deconvolutional layers as developed in DeeperCut [10]. As pose-estimation in the lab is typically simpler than most benchmarks in computer vision, we were able to remove aspects of DeeperCut that were not required to achieve human-level accuracy on several laboratory tasks [12]. This removal of the pairwise refinement,as well as our addition of integer linear programming on top of the part detectors significantly increased the inference speed (full multi-human DeeperCut: 578-1171 seconds per frame vs. 10-90 frames per second). Furthermore, we recently implemented faster inference for the feature detectors [27] (see below).

However, if a user aims to track (adult) human poses, many excellent options exist, including DeeperCut, ArtTrack, DeepPose, OpenPose, and OpenPose-Plus [2, 10, 11, 30–33, 35]. Some of these methods also allow real-time inference. To our knowledge, the performance of those methods has not been investigated on non-human animals. Conversely, since the trained networks can be used they provide excellent tools for pose-estimation of humans. There are also specific networks for faces [32] and hands [33]. However, if (additional) body parts beyond the body parts contained in the pre-trained networks need to be labeled, then DeepLabCut could also be useful.

While DeepLabCut, to the best of our knowledge, is the first example of using deep learning in the laboratory for pose-estimation, recently another deep learning package called LEAP was described [21]. LEAP requires the user to align the subject in the center of the frame, and there is some cost to rotations of the animal in terms of accuracy [21]. In contrast, DeepLabCut is robust for estimating the pose of animals that were not aligned. DeepLabCut was also shown to generalize to novel individuals (both flies and mice), as well as multiple mice, and conditions [12]. Certain behaviors are also not particularly amendable to alignment, like a Drosophila moving in 3D space [12].

DeepLabCut was shown to need less than 150 frames of human-labeled data for even challenging 3D hand movements of a mouse [12]. LEAP requires more training data to match similar performance, i.e. human-level accuracy. While the authors highlight an interactive training framework (which can be done with DeepLabCut if desired), and suggest starting with tens of frames, it needs approximately 500 images to achieve less than 3 pixel error on 192×192 pixel frames [21]. For example, DeepLabCut achieves < 5 pixel error with 100 labeled frames on a 800×800 pixel sized data set and reaches human level accuracy of 2.7 pixel with 500 labeled frames. Note this means the relative error (in relation to the frame size) is more than 4 times lower for DeepLabCut than for LEAP. These advantages seem to be due to transfer learning [12], while LEAP is trained from scratch.

Inference speed (i.e. the run time of the trained network on novel videos) may also be a consideration for users. While DeepLabCut takes 30-100 FPS (depending on the frame size) to process the data (if the batch size is 1). This means that it achieves real-time processing if the camera speed is less than 30-100 FPS. In contrast, LEAP may be faster, due to its 15 layer convolutional network, as the authors report 185 FPS with a batch size of 32 images for 192 x 192 sized images [21]. For similarly sized videos DeepLabCut runs at 90 FPS with batch size of 1 and accelerates substantially (up to 1000 FPS) for larger batch sizes [27].

## Advantages

DeepLabCut has been applied to a range of organisms with diverse visual background challenges. The main advantages are (i) our code guides the experimenter with a step-by-step procedure from labeling data to automated pose-extraction in a fast and efficient way; (ii) minimizes the cost of manual behavior analysis and with only a small number of training images can achieve human level accuracy; (iii) eliminates the need to put visible markers on the locations of interest to track the positions; (iv) can be easily adapted to analyze behaviors across species; (v) is open-source and free. Due to the use of deep features, DeepLabCut can learn to robustly extract body parts even with cluttered and varying background, inhomogeneous illumination, camera distortions, etc [12]. Thus, experimenter’s can perhaps design studies more around their scientific question, rather than constraints imposed by previous tracking algorithms. For example, it is common to image mice in a plain white, grey or black background to contrast heavily with their coat color. Now, even natural backgrounds, such as home cage bedding, natural grass, or dynamically changing backgrounds, as in trail-tracking [12], can be used. This is perhaps best illustrated by the cheetah example provided in this paper (Figures 1 and 7).

**FIG. 7.**
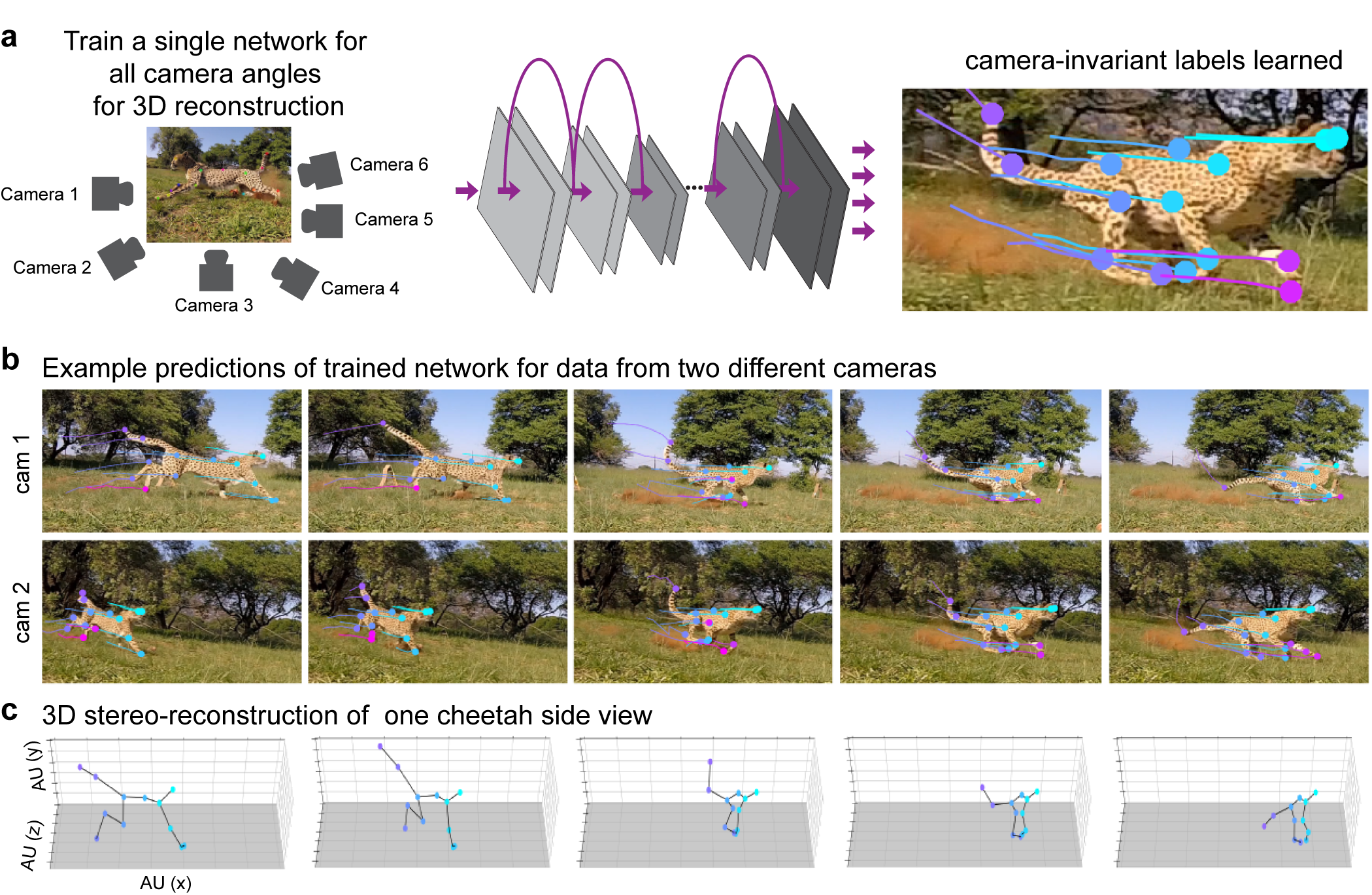
An example use case for pose estimation of a hunting cheetah. (a) Images collected from multiple cameras can be used to train a single network. (b) Example frames with overlaid body parts extracted by DeepLabCut, as well as a trajectory of past movements from previous frames. The same network was trained for multiple views, and is used to extract the poses. (b) 3D reconstruction of body parts from cheetah’s side facing both cameras from the same series of frames as in b.

DeepLabCut also allows for 3D pose estimation via multi-camera use. A user can easily a train network for each camera view and use standard camera calibration techniques [36] to resolve 3D locations. Alternatively, a user can combine multiple camera views into one network (i.e. see Figure 7 below and [27]), if the total pixel size is not too large given the available hardware, and then train this single network on the combined view. DeepLabCut does not require images to have a fixed frame size, as the feature detectors are not insensitive (within bounds) to the size due to automatic re-scaling during training [12]. There are also no camera requirements. Color and greyscale images captured from scientific cameras or consumer-grade cameras such as a GoPro can be used (and the package supports multiple video types such as .mp4 and .avi). Also we recently showed that DeepLabCut is robust to video-compression, potentially saving users *>*100 fold on data storage [27].

In addition to providing step-by-step procedures, our protocol allows the researcher to work on various projects simultaneously while keeping each of the projects isolated (on the computer). Moreover, the toolbox provides different frame-extraction methods to accommodate the analysis of videos from varying recording sessions. The toolbox is modular and by changing desired parameters at execution, the user can, for instance, utilize different frame extraction methods or identify frames which require further inspection. Reliably defining the user-defined labels across different frames is of paramount importance for supervised learning approaches like DeepLabCut. The labels can be body parts, or any other readily visible (part of an) object. The toolbox also provides interactive tools (graphical user interfaces, GUIs) to create, edit, and even add additional, new labels at a later stage of the project. The labels are stored in human readable and efficient data structures. The code also generates output files/directories for essential intermediate and final results in an automatic fashion. We believe that the error messages are intuitive, and enable researchers to efficiently utilize the toolbox. Furthermore, the toolbox facilitates visualization of the labeled data and creates videos with the extracted labels overlaid, making the entire toolbox a complete package for behavioral tracking. The extracted poses per video can also further analyzed with Python or imported to many other programs for further analysis.

## Limitations

One main limitation is that the toolbox requires modern computational hardware to produce fast and efficient results (namely, Graphical Processing Units -GPUs [5]). However, it is possible to run this toolbox on a standard computer with a compromise on the speed of analysis. It is about 10-100x slower [27]. However, the availability of inexpensive consumer-grade GPUs has opened the door for labs to perform advanced image processing in an autonomous way (i.e. in the lab, without lengthy transferring of data to centralized clusters).

Another consideration is that deep learning networks scale with the size of pixels, therefore larger images will be processed slower. Thus, for applications that would benefit from rapid tracking of the input images should be downsampled to achieve high rates (i.e. xs~90 frames per second (FPS) for a frame size of 138 × 138 with a batch size of 1). However, with new code updates and batch processing options, one can rapidly increase analysis speed. We recently demonstrated that with a batch size of 64, for a frame size of approx. 138 × 138 one can achieve 600 FPS [27].

Another limitation is that DeepLabCut, because it is designed to be general purpose, does not rely on heuristics such as a body model, and therefore occluded points cannot be tracked. However, DeepLabCut outputs a confidence score, which reports if a body part is actually visible. As we showed, one can (for instance) tell which legs are visible in a fly moving through a 3D space, or whether the finger tips of a reaching mouse are grasping the joystick and are thus occluded [12]. Furthermore, since DeepLabCut performs frame-by-frame prediction, even if due to occlusions, motion blur, etc, features cannot be detected in a few frames, they will be detected as soon as they are visible (unlike in many (actual) tracking methods, which required consistency across frames, such as the widely used Lucas —Kanade method [37]).

## Equipment Software

- Operating System: Linux (Ubuntu) or Windows (10). However, the authors recommend Ubuntu 16.04 LTS.
- DeepLabCut: The actively maintained toolbox is freely available at https://github.com/AlexEMG/ DeepLabCut. The code is written in Python 3 [38] and TensorFlow [39] for the feature detectors [10]. It can be simply installed with “pip install deeplabcut”, see the installation guide below.
- Anaconda: a free and open source distribution of the Python programming language (download from: https://www.anaconda.com/). DeepLabCut is written in Python 3 (https://www.python.org/) and not compatible with Python 2.
- TensorFlow [39]: an open-source software library for Deep Learning. The toolbox is tested with TensorFlow versions 1.0 to 1.4, 1.8, 1.10 and 1.11. Any one of these versions can be installed from https://www.tensorflow.org/install/
- OPTIONAL -Docker [40]: It is recommended to use the supplied docker container which includes a pre-installed TensorFlow with GPU support. This container builds on the nvidia-docker, which is only supported in Ubuntu.

## Hardware

- Computer: Any modern desktop workstation will be sufficient, as long as it has a PCI slot space as well as sufficient power supply for a GPU (see next item). The toolbox can also be used on laptops (e.g. for labeling data), then training can occur either on a CPU or elsewhere with a GPU. To note, training and evaluating the feature detectors will be slow without a GPU, but it is possible [27]. We recommend 32 GB of RAM on the system for CPU for analysis, but this is not a hard minimum. More information on optimally running TensorFlow can be found here: https://www.tensorflow.org/guide/performance/overview.
- GPU: It is recommended to have a GPU with at least 8 GB memory such as the NVIDIA GeForce 1080. However, is it not required to use a GPU, a CPU is sufficient but training and evaluating the network steps are considerably slower [27]. This toolbox can also be used on cloud computing services (such as Google Cloud/Amazon Web Services).
- Camera(s): The toolbox is robust to extraction of poses from data collected by many cameras. There are no *a priori* limitations in terms of lighting; color or greyscale images are acceptable, videos captured under infrared light, and inhomogeneous or natural lighting can be used. Cameras should be placed such that the features the user wishes to track are visible. For reference, as reported in [12], the following cameras have been used for video capture: Point Grey Firefly (FMVU-03MTM-CS), Grasshopper 3 4.1MP Mono USB3 Vision (CMOSIS CMV4000-3E12), infrared-sensitive CMOS camera from Basler. Here, the Cheetahs were filmed with GoPro Hero 5 Session cameras, and DFK-37BUX287 cameras from The Imaging Source, Inc. were used to film mice.

## Installation

**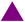CRITICAL POINT**: We recommend that the user first simply installs DeepLabCut in an Anaconda environment, as they can access all steps on a CPU. We also provide Jupyter Notebooks for quick demonstrations of the workflow.

Then for GPU support, the user can either set up TensforFlow, CUDA and the NVIDIA card driver on their own computer (following NVIDIA and TensorFlow’s operation system and graphic card specific instructions provided, see below for links), or alternatively, they can use our supplied Docker container that has TensorFlow and CUDA pre-installed.

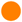**Timing:** It takes around 10-60 minutes to install the toolbox, depending on the installation method chosen.

All the following commands will be run in the app “terminal” (in Ubuntu, and called “cmd” in Windows). Please first open the terminal (search “terminal” or “cmd”).

**Warning:** By running “pip install deeplabcut”, all required distributions are installed (except wxPython and TensorFlow). If you perform a system wide installation, and the computer has other Python packages or TensorFlow versions installed that conflict, this will overwrite them. If you have a dedicated machine for DeepLabCut, this is fine. If there are other applications that require different versions of libraries, then one would potentially break those applications. One solution to this problem is to create a virtual environment, a self-contained directory that contains a Python installation for a particular version of Python, plus additional packages. One way to manage virtual environments is to use conda environments (for which you need Anaconda installed).

An Anaconda environment can be created in the following way:

~~~
conda create -n <name of the environment> python=3.6
~~~

This environment can be accessed by typing:

~~~
in Windows: activate <name of the environment>
in Linux: source activate <name of the environment>
~~~

Once the environment was activated, the user can install DeepLabCut, as described below.

A user can exit the conda environment at any time by typing the following command in the terminal:

~~~
in Windows: deactivate <name of the environment>
in Linux: source deactivate <name of the environment>
~~~

The user can re-activate the environment as shown above. The toolbox is installed within the environment and once the user deactivates it, they must re-enter the environment use the DeepLabCut toolbox.

**To install DeepLabCut**, in the terminal type:

pip install deeplabcut

For GUI support in the terminal type:

~~~
Windows: pip install -U wxPython
Linux (Ubuntu 16.04): pip install https://extras.wxpython.org/wxPython4/extras/linux/gtk3/ubuntu-16.04/wxPython-4.0.3-cp36-cp36m-linux_x86_64.whl
~~~

As users may vary in their use of a GPU or CPU, TensorFlow is not installed with the command “pip install deeplabcut”. For CPU only support, in the terminal please type:

~~~
pip install tensorflow==1.10
~~~

If you have a GPU, you should then install the NVIDIA CUDA package and an appropriate driver for your specific GPU. Please follow the instructions found here, https://www.tensorflow.org/install/gpu, as we cannot cover all the possible combinations that exist (and these are continually updated packages).

**DeepLabCut Docker:** When using a GPU, it is recommended to use the supplied Docker container if you use Ubuntu, as TensorFlow and DeepLabCut are already installed. Unfortunately, a dependency (nvidia-docker) is currently not compatible with Windows. Additionally, the GUIs cannot be used inside the Docker container. We envision a user installing DeepLabCut into an Anaconda environment, but then executing the GPU steps inside the container. This way, installation is extremely simple. Full details on installing Docker4DeepLabCut2.0 can be found at: https://github.com/MMathisLab/Docker4DeepLabCut2.0.

Lastly, it is also possible to run the network training steps of DeepLabCut in the supplied Docker, or on cloud computing resources (such as AWS, Google, or on a University cluster). For running DeepLabCut on certain platforms you will need to suppress the GUI support. This can be done by setting an environment variable before loading the toolbox:

~~~
Linux: export DLClight=True
Windows: set DLClight=True
~~~

If you want to re-engage the GUIs (and have installed wxPython as described above):

~~~
Linux: unset DLClight
Windows: set DLClight=
~~~

## STEP-BY-STEP PROCEDURE

The following commands will guide the user on how to use the toolbox in python/ipython. We also provide an overview guide for all the commands in Step N, and provide Jupyter Notebooks, described in Section M.

For any DeepLabCut function, there is a ‘help’ document that defines all the required and optional functions. In ipython, simply place a ? after the function, i.e. deeplabcut.create_new_project?. See more options in Section O. To begin, open an ipython session and import the package by typing in the terminal:

~~~
ipython
import deeplabcut
~~~

## A. Create a New Project

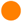**Timing**: The time required to create a new project 2 minutes.

The **function create_new_project** creates a new project directory, required subdirectories, and a basic project configuration file. Each project is identified by the name of the project (e.g. Reaching), name of the experimenter (e.g. YourName), as well as the date at creation. Thus, this function requires the user to input the enter the name of the project, the name of the experimenter, and the full path of the videos that are (initially) used to create the training dataset (without spaces in each, i.e. Test1 vs. Test 1).

Optional arguments specify the working directory, where the project directory will be created, and if the user wants to copy the videos (to the project directory). If the optional argument working_directory is unspecified, the project directory is created in the current working directory, and if copy_videos is unspecified symbolic links for the videos are created in the videos directory. Each symbolic link creates a reference to a video and thus eliminates the need to copy the entire video to the video directory (if the videos remain at that original location).

~~~
>> deeplabcut.create_new_project(‘Name of the project’,‘Name of the experimenter’, [‘Full path of video 1’,‘Full path of video2’,‘Full path of video3’], working_directory=‘Full path of the working directory’,copy_videos=True/False)
~~~

To automatically create a variable that points to the ‘config_path’ variable, which will be used below, add:

~~~
>> config_path = deeplabcut.create_new_project(...
~~~

This set of arguments will create a project directory with the name **“Name of the project+name of the experimenter+date of creation of the project”** in the “Working directory” and creates the symbolic links to videos in the “videos” directory. The project directory will have subdirectories: “dlc-models”, “labeled-data”, “training-datasets”, and “videos”. All the outputs generated during the course of a project will be stored in one of these subdirectories, thus allowing each project to be curated in separation from other projects. The purpose of the subdirectories is as follows:

- **dlc-models:** This directory contains the subdirectories “test” and “train”, each of which holds the meta information with regard to the parameters of the feature detectors in configuration file. The configuration files are YAML files, a common human-readable data serialization language. These files can be opened and edited with standard text editors. The subdirectory “train” will store checkpoints (called snapshots in TensorFlow) during training of the model. These snapshots allow the user to reload the trained model without re-training it, or to pick-up training from a particular saved checkpoint, in case the training was interrupted.
- **labeled-data**: This directory will store the frames used to create the training dataset. Frames from different videos are stored in separate subdirectories. Each frame has a filename related to the temporal index within the corresponding video, which allows the user to trace every frame back to its origin.
- **training-datasets**: This directory will contain the training dataset used to train the network and metadata, which contains information about how the training dataset was created.
- **videos**: Directory of video links or videos. When copy_videos is set to “False”, this directory contains symbolic links to the videos. If it is set to “True” then the videos will be copied to this directory. The default is “False”. Additionally, if the user wants to add new videos to the project at any stage, the function add_new_videos can be used. This will update the list of videos in the project’s configuration file.

~~~
>> deeplabcut.add_new_videos(‘Full path of the project configuration file*’,[‘full path of video 4’, ‘full path of video 5’],copy_videos=True/False)
~~~

*Please note, ‘Full path of the project configuration file’ will be referenced as a variable called config_path throughout this protocol.

The project directory also contains the main configuration file called *config.yaml*. The *config.yaml* file contains many important parameters of the project. A complete list of parameters including their description can be found in Box 1.

The ‘create a new project’ step writes the following parameters to the configuration file: *Task*, *scorer*, *date*, *project_path* as well as a list of videos *video_sets*. The first three parameters should not be changed. The list of videos can be changed by adding new videos or manually removing videos.

### Box 1: Glossary of parameters in the project configuration file (config.yaml)

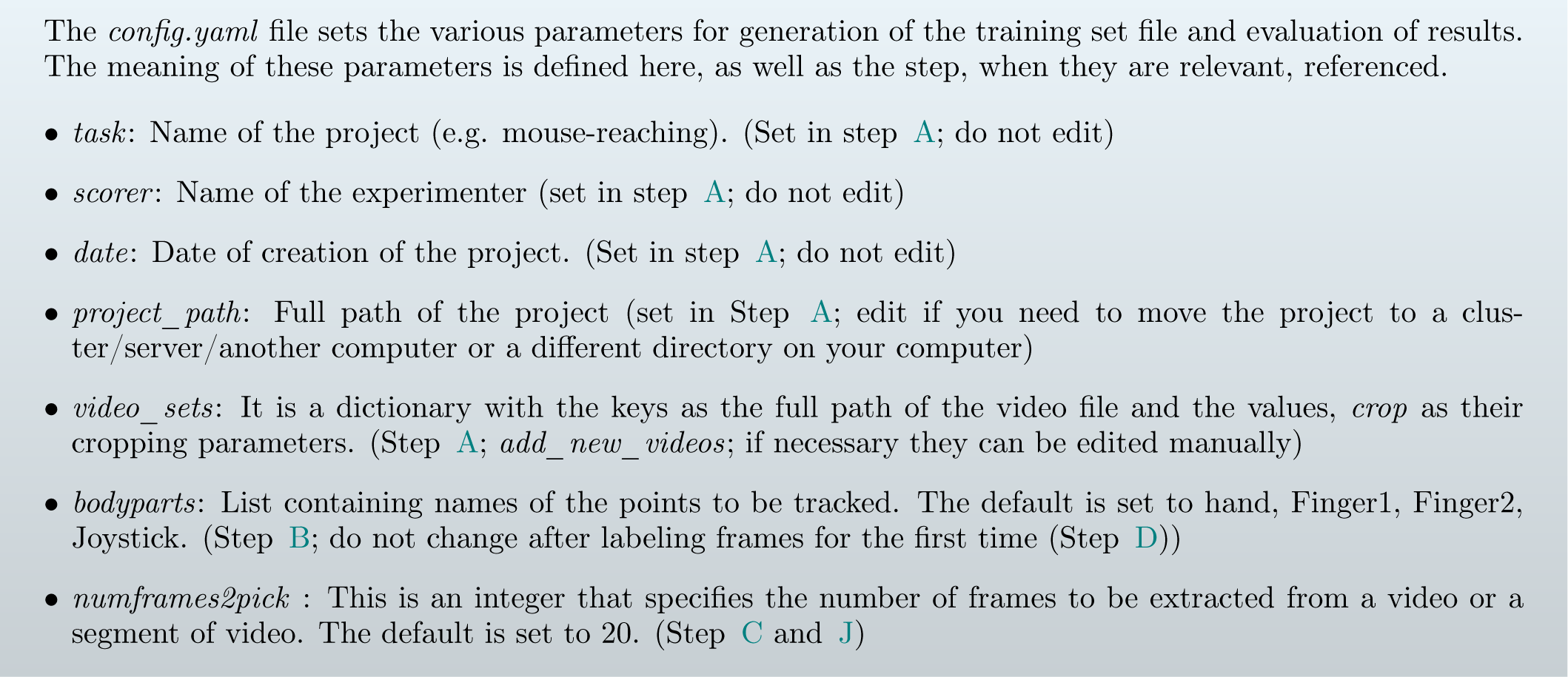
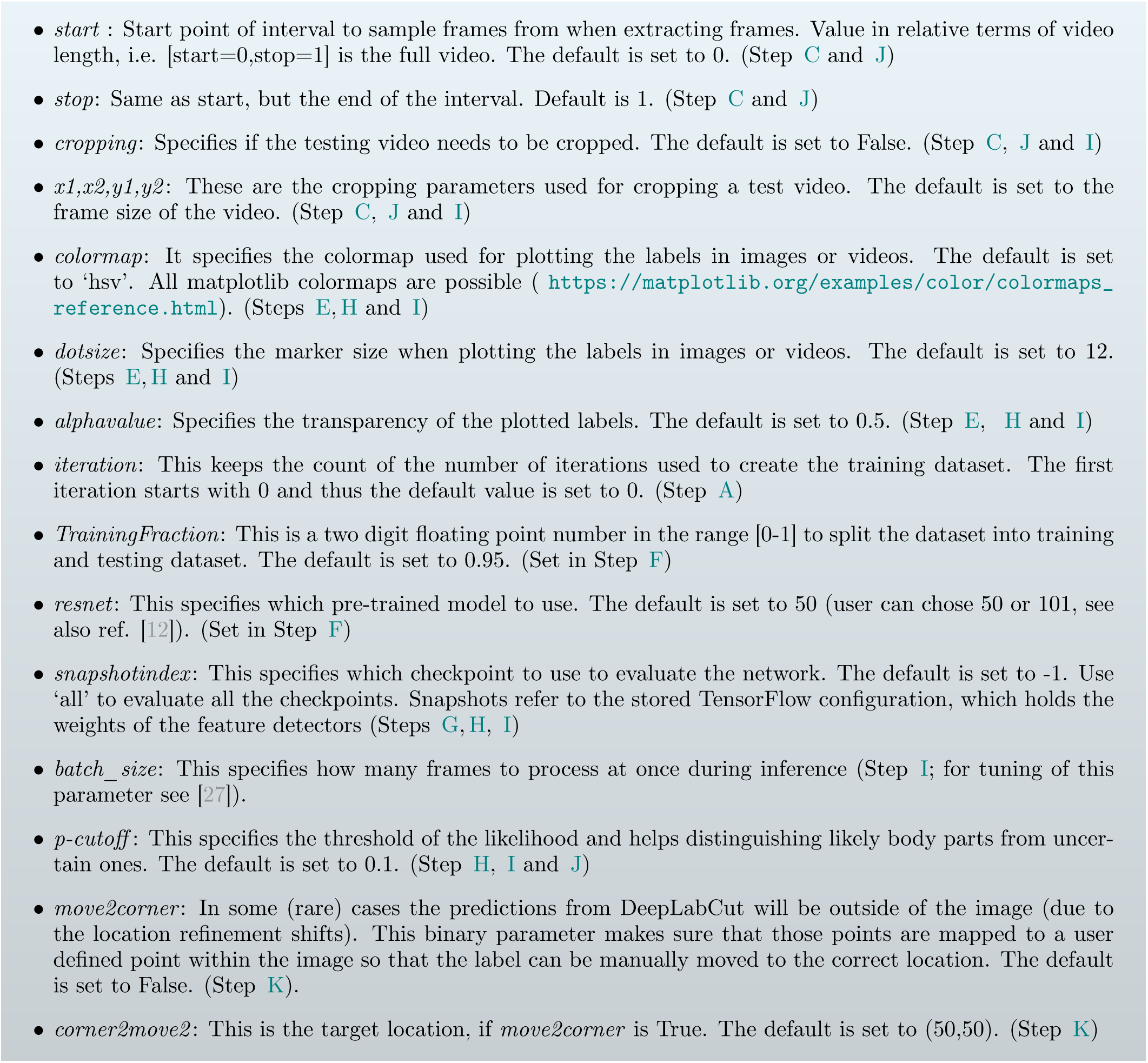

## B. Configure the Project

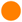**Timing:** The time required to edit the *config.yaml* file is *≈* 5 minutes.

Next, open the *config.yaml* file, which was created during create_new_project. You can edit this file in any text editor. Familiarize yourself with the meaning of the parameters (Box 1). You can edit various parameters, in particular add the list of *bodyparts* (or points of interest) that you want to track. For the next data selection step *numframes2pick*, *start*, *stop*, *x1, x2, y1, y2* and *cropping* are of major importance.

## C. Data Selection

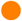**Timing:** The time required to select the data is mainly related to the mode of selection, number of videos, and the algorithm used to extract the frames (Figure 3). It takes ≈ 30 sec/video in an automated mode with default parameter settings (processing time strongly depends on length and size (seconds) of video). However, it may take longer in the manual mode and mainly depends on the time it takes to find frames with diverse behavior.

**FIG. 3.**
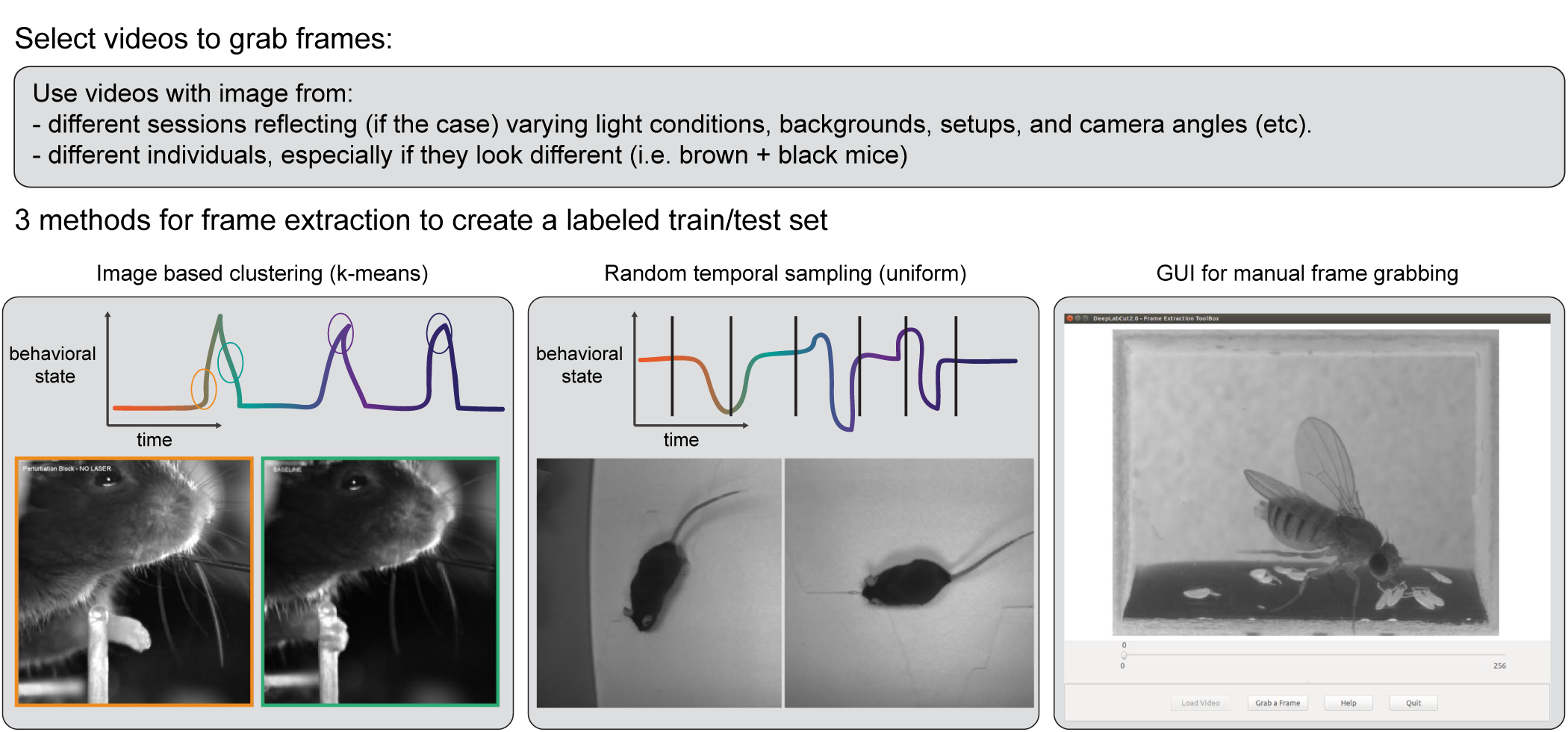
Three methods for frame selection. The toolbox contains three methods to extracting frames, namely, by clustering based on visual content, by randomly sampling in uniform way across time, or by manually grabbing frames of interest using a custom GUI. Depending on the studied behavior, the appropriate method can be used. Additional methods will be added in the future.

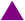**CRITICAL POINT:** A good training dataset should consist of a sufficient number of frames that capture the full breadth of the behavior. This implies to select the frames from different (behavioral) sessions and different animals, if those vary substantially (to train an invariant, robust feature detector). Thus, a good training dataset should reflect the diversity of the behavior with respect to postures, luminance conditions, background conditions, animal identities, etc. of the data that will be analyzed. For the behaviors we have tested so far, a data set of 100−200 frames gave good results [12]. However, depending on the required accuracy and the nature of the scene statistics, more or less frames might be necessary to create the training data set. Ultimately, in order to scale up the analysis to large collections of videos with perhaps unexpected conditions, one can also refine the data set in an adaptive way (Step J and Step K).

The function *extract_frames* extracts the random frames from all the videos in the project configuration file in order to create a training dataset. The extracted frames from all the videos are stored in a separate subdirectory named after the video file’s name under the ‘labeled-data’. This function also has various parameters that might be useful based on the user’s need. The default values are ‘automatic’ and ‘kmeans’.

~~~
>> deeplabcut.extract_frames(config_path,‘automatic/manual’,‘uniform/kmeans’)
~~~

**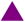CRITICAL POINT:** It is advisable to keep the frame size small, as large frames increase the training and inference time. The cropping parameters for each video can be provided in the *config.yaml* file (and see below).

When running the function *extract_frames*, if the parameter *crop*=True and *checkcropping* =True, then it will crop the frames to the size provided in the *config.yaml* file, and the user can first check the bounding box of the cropping. Upon calling *extract_frames* a image will pop up with a red bounding box based on the crop parameters so that the user can check those parameters. Once the user closes the pop-up window, they will be asked if the cropping is correct. If yes, then the frames are extracted accordingly. If not, the cropping parameters can be iteratively adjusted based on this graphical feedback before proceeding. As as reminder, for each function, place a ‘ ?’ after the function (i.e. deeplabcut.extract_frames?) to see all the available options.

The provided function either selects frames from the videos in a randomly and temporally uniformly distributed way (uniform), by clustering based on visual appearance (k-means), or by manual selection (Figure 3). Random selection of frames works best for behaviors where the postures vary across the whole video. However, some behaviors

## Select videos to grab frames

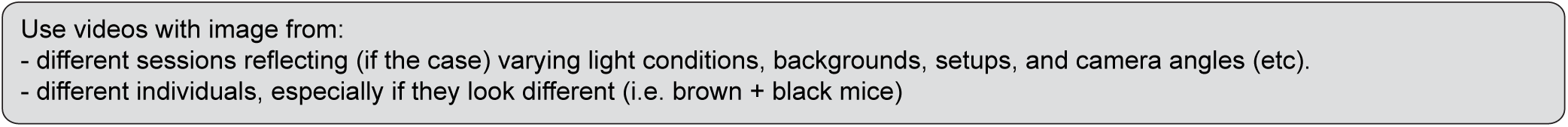

3 methods for frame extraction to create a labeled train/test set might be sparse, as in the case of reaching where the reach and pull are very fast and the mouse is not moving much between trials. In such a case, the function that allows selecting frames based on k-means derived quantization would be useful. If the user chooses to use k-means as a method to cluster the frames, then this function downsamples the video and clusters the frames using k-means, where each frame is treated as a vector. Frames from different clusters are then selected. This procedure makes sure that the frames look different. However, on large and long videos, this code is slow due to computational complexity.

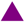**CRITICAL POINT**: It is advisable to extract frames from a period of the video that contains interesting behaviors, and not extract the frames across the whole video. This can be achieved by using the *start* and *stop* parameters in the *config.yaml* file. Also, the user can change the number of frames to extract from each video using the *numframes2extract* in the *config.yaml* file.

However, picking frames is highly dependent on the data and the behavior being studied. Therefore, it is hard to provide all purpose code that extracts frames to create a good training dataset for every behavior and animal. If the user feels specific frames are lacking, they can extract hand selected frames of interest using the interactive GUI provided along with the toolbox. This can be launched by using:

~~~
>> deeplabcut.extract_frames(config_path,‘manual’)
~~~

The user can use the ‘Load Video’ button to load one of the videos in the project configuration file, use the scroll bar to navigate across the video and ‘Grab a Frame’ to extract the frame. The user can also look at the extracted frames and e.g. delete frames (from the directory) that are too similar before re-loading the set and then manually annotating them.

## D. Label Frames

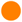**Timing**: The time required to label the frames depends on the speed of the experimenter to identify the correct label and is mainly related to the number of body parts and the total number of extracted frames.

The toolbox provides a function ‘label_frames’ which helps the user to easily label all the extracted frames using an interactive graphical user interface (GUI). The user should have already named the body parts to label (points of interest) in the project’s configuration file by providing a list. The following command invokes the labeling toolbox.

~~~
>> deeplabcut.label_frames(config_path)
~~~

The GUI is launched taking the size of the user’s screen into account, so if two monitors are used (in landscape mode) place ‘Screens=2’ after config_path (The default is 1). For other configurations, please see the linked troubleshooting guide, Section (O).

Next, the user needs to use the ‘Load Frames’ button to select the directory which stores the extracted frames from one of the videos. A right click places the first body part, and subsequently, the user can either select one of the radio buttons (top right) to select a body part to label, or there is a built in auto-advance to the next body part. If a body part is not visible, simply do not label the part and click on the next body part you can label. Clicking the right arrow key will advance to the next frame. Each label will be plotted as a dot in a unique color (see Figure 4 for more details).

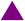**CRITICAL POINT**: Finalize the position of the selected label before changing the dot size for the next labels. it is recommended to do this once after the first frame is labeled. Then, proceed to the next frame.

**FIG. 4.**
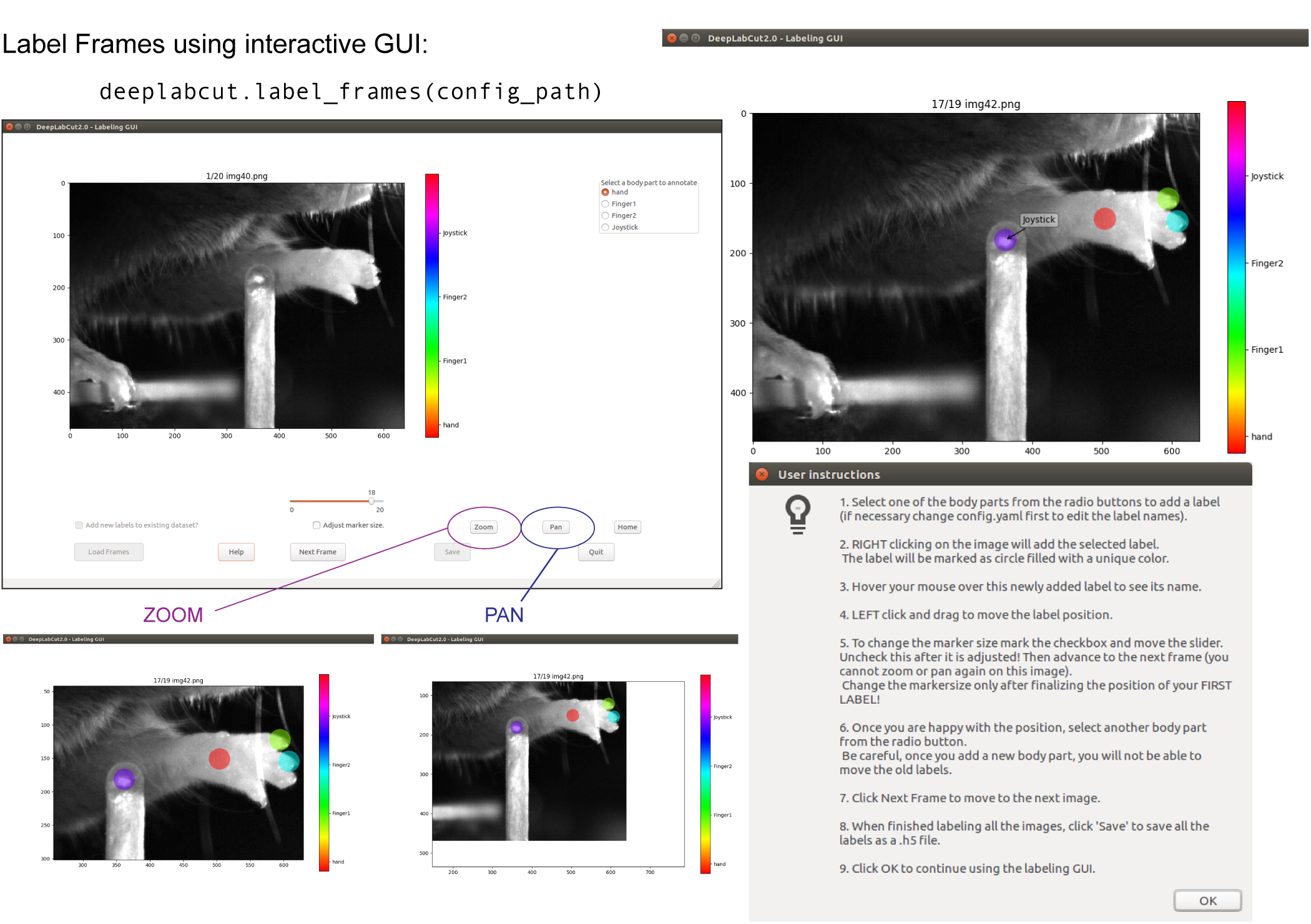
Labeling GUI: The toolbox contains a labeling GUI that allows for frame loading, labeling, re-adjustments, and saving the dataset into the correct format for future steps. An example label is shown, and the help functions are described. Additionally, a user can decide to add more points to an existing labeled dataset by adding the new labels to the *config.yaml* file and re-opening the labeling GUI and then the user can click the tick box in the bottom left corner to add new points.

The user is free to move around the body part with a left click and drag once they satisfied with its position, can select another radio button (in the top right) to switch to the respective body part. Once the user starts labeling a subsequent body part, preceding labels of the body parts can no longer be moved. The user can skip a body part if it is not visible. Once all the visible body parts are labeled, then the user can use ‘Next Frame’ to load the following frame. The user needs to save the labels after all the frames from one of the videos are labeled by clicking the save button at the bottom right. Saving the labels will create a labeled dataset for each video in a hierarchical data file format (HDF) in the subdirectory corresponding to the particular video in ‘labeled-data’.

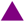**CRITICAL POINT**: It is advisable to consistently label similar spots (e.g. on a wrist that is very large, try to label the same location). In general, invisible or occluded points should not be labeled by the user. They can simply be skipped by not applying the label anywhere on the frame.

OPTIONAL: In an event of adding more labels to the existing labeled dataset, the user need to append the new labels to the *bodyparts* in the *config.yaml* file. Thereafter, the user can call the function ‘label_frames’ and check the left lower tick box, ‘Add new labels to existing dataset?’ before loading the frames. Saving the labels after all the images are labelled will append the new labels to the existing labeled dataset.

## E. Check Annotated Frames

**OPTIONAL**: Checking if the labels were created and stored correctly is beneficial for training, since labeling is one of the most critical parts for creating the training dataset. The DeepLabCut toolbox provides a function ‘check_labels’ to do so. It is used as follows:

~~~
>> deeplabcut.check_labels(config_path)
~~~

For each video directory in *labeled-data* this function creates a subdirectory with ‘labeled’ as a suffix. Those directories contain the frames plotted with the annotated body parts. The user can double check if the body parts are labeled correctly. If they are not correct, the user can call the refinement GUI (Step K and check the tick box for “adjust original labels" to adjust the location of the labels).

## F. Create Training Dataset

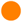**Timing**: The time required to create a training dataset is <1 minute. Combining the labeled datasets from all the videos and splitting them will create train and test datasets. The training data will be used to train the network, while the test data set will be used for evaluating the network. The function ‘create_training_dataset’ performs those steps.

~~~
>> deeplabcut.create_training_dataset(config_path,num_shuffles=1)
~~~

The set of arguments in the function will shuffle the combined labeled dataset and split it to create train and test sets. The subdirectory with suffix ‘iteration#’ under the directory ‘training-datasets’ stores the dataset and meta information, where the ‘#’ is the value of ‘iteration’ variable stored in the project’s configuration file (this number keeps track of how often the dataset was refined).

**OPTIONAL**: If the user wishes to benchmark the performance of the DeepLabCut, they can create multiple training datasets by specifying an integer value to the *num_shuffles*.

Each iteration of the creation of a training dataset, will create a ‘.mat’ file, which is used by the feature detectors and a ‘.pickle’ file which contains the meta information about the training dataset. This also creates two subdirectories within ‘dlc-models’ called ‘test’ and ‘train’, and these each have a configuration file called *pose_cfg.yaml*. Specifically, the user can edit the *pose_cfg.yaml* within the train subdirectory before starting the training. These configuration files contain meta information with regard to the parameters of the feature detectors. Key parameters are listed in Box 2.

### Box 2: Parameters of interest in the configuration file, pose_cfg.yaml.

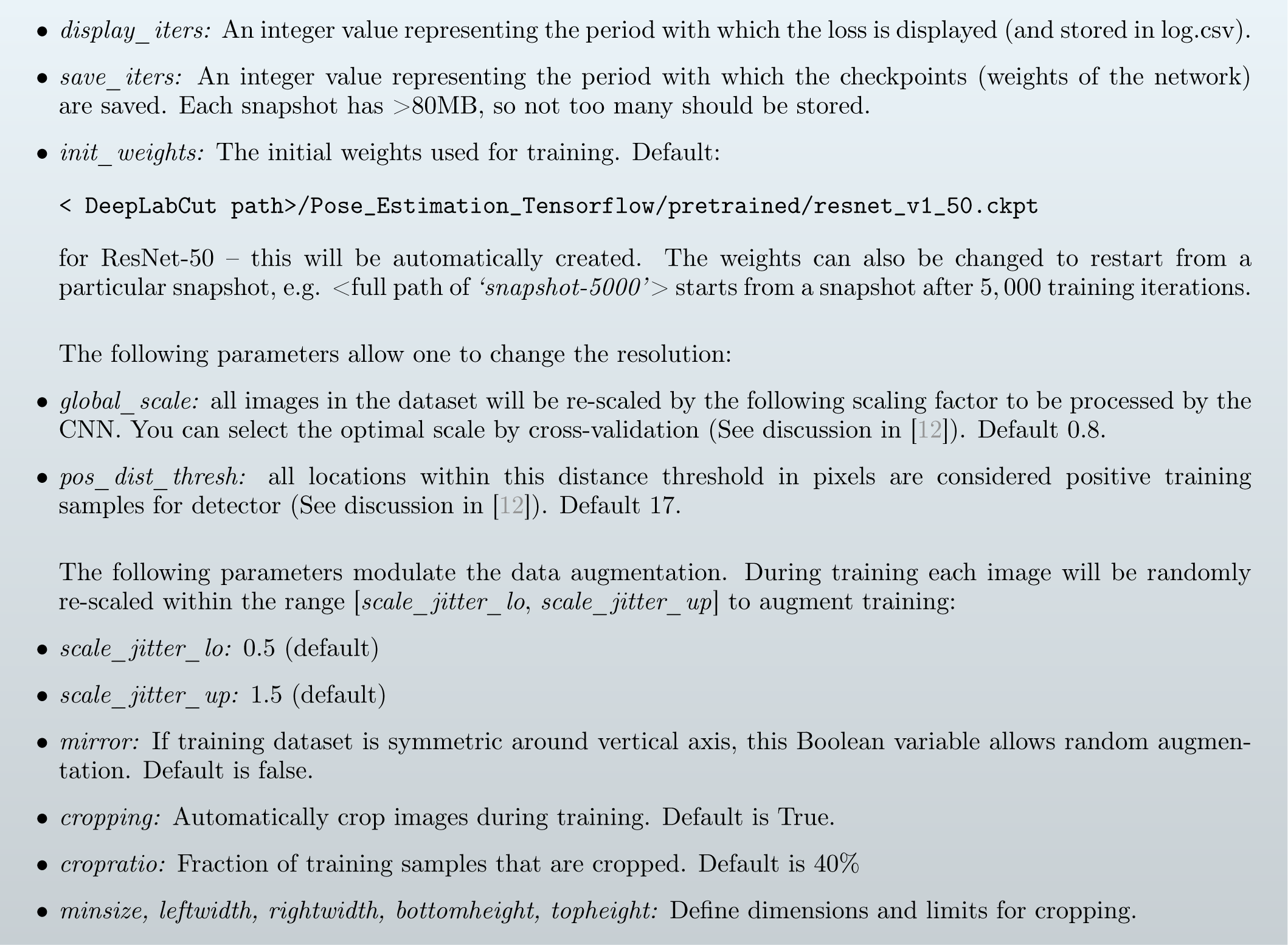

## G. Train The Network

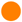**Timing**: The time required to train the network mainly depends on the frame size of the dataset and the computer hardware. On a NVIDIA GeForce GTX 1080 Ti GPU, it takes ≈ 6 hrs to train the network for at least 200,000 iterations. On the CPU, it will take several days to train for the same number of iterations on the same training dataset.

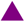**CRITICAL POINT**: It is recommended to train for thousands of iterations until the loss plateaus (typically *>*200,000). The variables ‘display_iters’ and ‘save_iters’ in the *pose_cfg.yaml* file allows the user to alter how often the loss is displayed and how often the weights are stored.

The function ‘train_network’ helps the user in training the network. It is used as follows:

~~~
>> deeplabcut.train_network(config_path,shuffle=1)
~~~

The set of arguments in the function starts training the network for the dataset created for one specific shuffle. Example parameters that one can call:

~~~
train_network(config_path,shuffle=1,trainingsetindex=0,gputouse=None,max_snapshots_to_keep=5, autotune=False,displayiters=None,saveiters=None)
~~~

Important Parameters:

config: Full path of the config.yaml file as a string.

shuffle: Integer value specifying the shuffle index to select for training. Default is set to 1

trainingsetindex: Integer specifying which Training set Fraction to use. By default the first (note that Training Fraction is a list in config.yaml).

gputouse: Natural number indicating the number of your GPU (see number in nvidia-smi). If you do not have a GPU put None. see also: https://nvidia.custhelp.com/app/answers/detail/a_id/3751/~/useful-nvidia-smi-queries

Additional parameters:

max_snapshots_to_keep: This sets how many snapshots are kept, i.e. states of the trained network. Every saving iteration a snapshot is stored, however only the last max_snapshots_to_keep many are kept! If you change this to None, then all are kept. See also: https://github.com/AlexEMG/DeepLabCut/issues/8#issuecomment-387404835 autotune: property of TensorFlow, tested to be faster if set to ‘False’; see https://github.com/tensorflow/tensorflow/issues/13317). Default: False displayiters: this variable is actually set in pose_config.yaml. However, you can overwrite it with this. If None, the value from pose_config.yaml is used, otherwise it is overwritten! Default: None saveiters: this variable is set in pose_config.yaml. However, you can overwrite it with this. If None, the value from there is used, otherwise it is overwritten! Default: None By default, the pre-trained ResNet network is not provided in the DeepLabCut toolbox (as it has around 100MB). However, if not previously downloaded from the TensorFlow model weights, it will be downloaded and stored in a subdirectory ‘pre-trained’ under the subdirectory ‘models’ in ‘Pose_Estimation_Tensorflow’, where deeplabcut is installed (i.e. usually site-packages). At user specified iterations during training checkpoints are stored in the subdirectory ‘train’ under the respective iteration directory. If the user wishes to restart the training at a specific checkpoint they can specify the full path of the checkpoint to the variable ‘init_weights’ in the *pose_cfg.yaml* file under the ‘train’ subdirectory (see Box 2).

## H. Evaluate the Trained Network

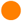**Timing**: The time required to evaluate the trained network depends on the computational hardware, number of checkpoints to evaluate, and the number of images in the training dataset. With the default parameters and around 150 images in the training dataset, it will take *≈* 2 mins/checkpoint on a GPU. However, for the same snapshot and training set a CPU computes *≈* ten times longer (i.e. 20 min).

It is important to evaluate the performance of the trained network. This performance is measured by computing the mean average Euclidean error (MAE; which is proportional to the average root mean square error) between the manual labels and the ones predicted by DeepLabCut. The MAE is saved as a comma separated file and displayed for all pairs and only likely pairs (*>*p-cutoff). This helps to exclude, for example, occluded body parts. One of the strengths of DeepLabCut is that due to the probabilistic output of the scoremap, it can, if sufficiently trained, also reliably report if a body part is visible in a given frame. (see discussions of finger tips in reaching and the Drosophila legs during 3D behavior in [12]). The evaluation results are computed by typing:

~~~
>> deeplabcut.evaluate_network(config_path,shuffle=[1], plotting=True)
~~~

Setting ‘plotting’ to true plots all the testing and training frames with the manual and predicted labels. The user should visually check the labeled test (and training) images that are created in the ‘evaluation-results’ directory. Ideally, DeepLabCut labeled unseen (test images) according to the user’s required accuracy, and the average train and test errors are comparable (good generalization). What (numerically) comprises an acceptable MAE depends on many factors (including the size of the tracked body parts, the labeling variability, etc.). Note that the test error can also be larger than the training error due to human variability (in labeling, see Figure 2 in [12]).

The plots can be customized by editing the *config.yaml* file (i.e. the colormap, scale, marker size (dotsize), and transparency of labels (alphavalue) can be modified). By default each body part is plotted in a different color (governed by the colormap) and the plot labels indicate their source. Note that by default the human labels are plotted as plus (‘+’), DeepLabCut’s predictions either as ‘.’ (for confident predictions with likelihood > p-cutoff) and ‘x’ for (likelihood *<*= p-cutoff). Examples test and training plots from various projects are depicted in Figure 5.

**FIG. 5.**
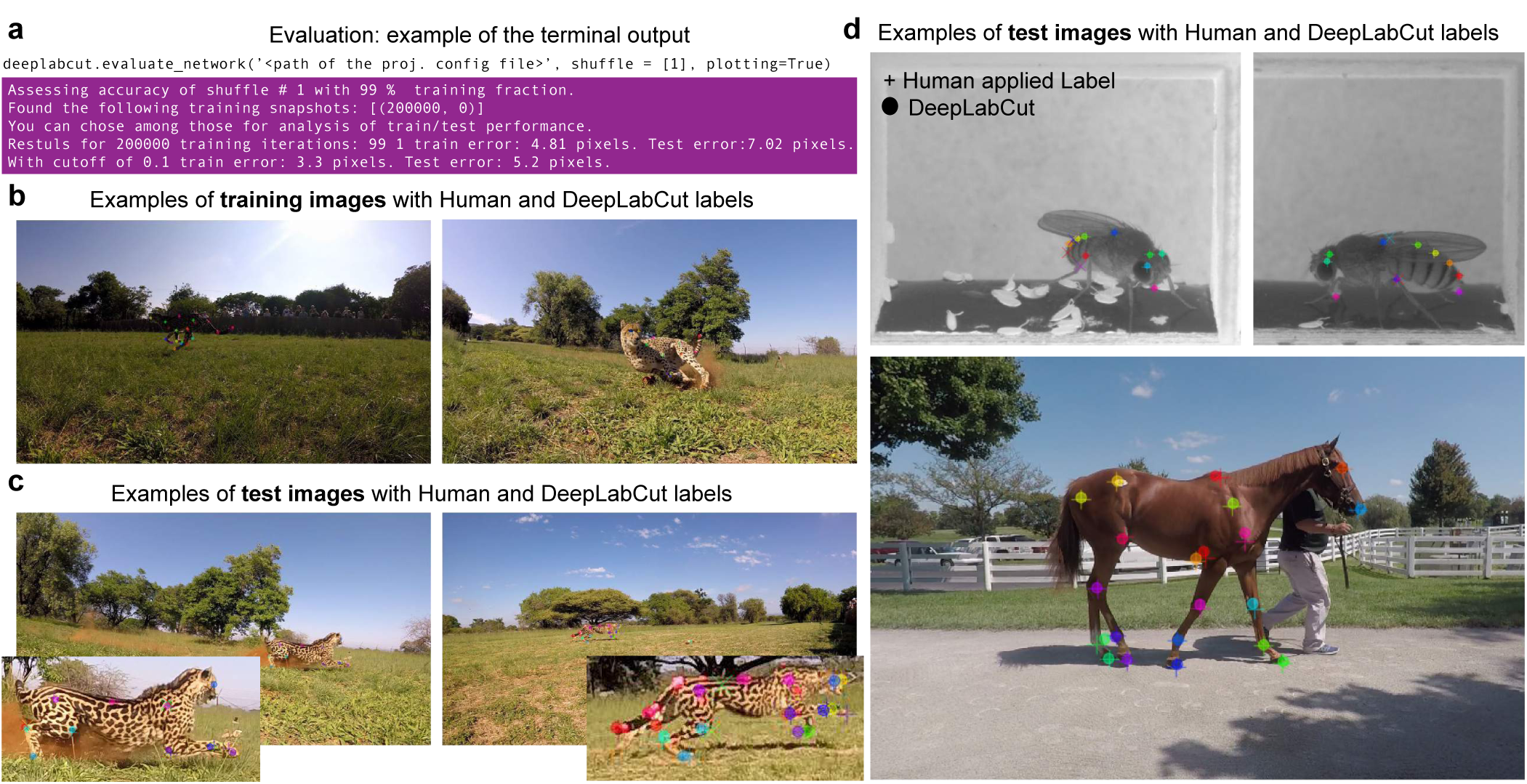
Evaluation results: (a) The code that is used to evaluate the network, and the output the user will see in the terminal. (b)Another output is labeled frames from training and (c) test images, as shown for a Cheetah project. (d) Additional example test evaluation images from Drosophila (different images from network trained in [12]) and horse tracking projects.

The evaluation results for each shuffle of the training dataset are stored in a unique subdirectory in a newly created directory ‘evaluation-results’ in the project directory. The user can visually inspect if the distance between the labeled and the predicted body parts is acceptable. In the event of benchmarking with different shuffles of same training dataset, the user can provide multiple shuffle indices to evaluate the corresponding network. If the generalization is not sufficient, the user might want to:

- check if the labels were imported correctly, i.e. invisible points are not labeled and the points of interest are labeled accurately (see Step E)
- make sure that the loss has already converged (Step G)
- consider labeling additional images and make another iteration of the training data set (Section J).

## I. Video Analysis and Plotting Results

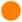**Timing**: The time required to analyze a video is mainly related to the number of frames in the video and if the computational power of a GPU is available. With a GPU analysis runs at 10-100 FPS depending on the frame size (for 1000 *×* 1000 to 200 *×* 200, respectively).

The trained network can be used to analyze new videos. The user needs to first choose a checkpoint with the best evaluation results for analyzing the videos. In this case, the user can enter the corresponding index of the checkpoint to the variable *snapshotindex* in the *config.yaml* file. By default, the most recent checkpoint (i.e. last) is used for analyzing the video. Then, a new video can be analyzed by typing:

~~~
>> deeplabcut.analyze_videos(config_path,[‘/analysis/project/videos/reachingvideo1.avi’], shuffle=1, save_as_csv=True, videotype=.avi)
~~~

The labels are stored in a MultiIndex Pandas Array ([41], http://pandas.pydata.org), which contains the name of the network, body part name, (*x, y*) label position in pixels, and the likelihood for each frame per body part. These arrays are stored in an efficient Hierarchical Data Format (HDF) in the same directory, where the video is stored. However, if the flag *save_as_csv* is set to True, the data can also be exported in comma-separated values format (*.csv*), which in turn can be imported in many programs, such as MATLAB, R, Prism, etc.; This flag is set to False by default.

The labels for each body part across the video (‘trajectories’) can also be plotted after analyze_videos is run. The provided plotting function in this toolbox utilizes matplotlib [42] therefore these plots can easily be customized by the end user. This function can be called by typing:

~~~
>>> deeplabcut.plot_trajectories(config_path,[‘/analysis/project/videos/reachingvideo1.avi’])
~~~

The trajectories can also be easily imported into many programs for further behavioral analysis. For example, a user can compute general movement statistics (position, velocity, etc), or interactions e.g. with (tracked) objects in the environment (i.e. with DeepLabCut there is no centering of the animal(s) therefore the user can easily measure relative movements within the background environment). The outputs from DeepLabCut can also interfaced with behavioral clustering tools such as JAABA [43], MotionMapper [44], MoSeq [45], or other clustering approaches such as iterative denoising tree (IDT) methods [46, 47], and more [48]. Indeed, users are contributing code for analysis of the outputs of DeepLabCut here: https://github.com/AlexEMG/DLCutils.

Additionally, the toolbox provides a function to create labeled videos based on the extracted poses by plotting the labels on top of the frame and creating a video. One can use it as follows to create multiple labeled videos:

~~~
>> deeplabcut.create_labeled_video(config_path,[‘/analysis/project/videos/reachingvideo1.avi’,‘/analysis/project/videos/reachingvideo2.avi’])
~~~

This function has various parameters, in particular the user can set the *colormap*, the *dotsize*, and *alphavalue* of the labels in *config.yaml* file.

## J. Refinement: Extract Outlier Frames

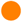**Timing**: The time required to select the data is mainly related to the mode of selection, number of videos and the algorithm used to extract the frames. It takes around 1 min/video in an automated mode with default parameter settings (processing time strongly depends on length and size of video).

While DeepLabCut typically generalizes well across datasets, one might want to optimize its performance in various, perhaps unexpected, situations. For generalization to large data sets, images with insufficient labeling performance can be extracted, manually corrected by adjusting the labels to increase the training set and iteratively improve the feature detectors. Such an active learning framework can be used to achieve a predefined level of confidence for all images with minimal labeling cost (discussed in [12]). Then, due to the large capacity of the neural network that underlies the feature detectors, one can continue training the network with these additional examples. One does not necessarily need to correct all errors as common errors could be eliminated by relabeling a few examples and then re-training. A priori, given that there is no ground truth data for analyzed videos, it is challenging to find putative “outlier frames”. However, one can use heuristics such as the continuity of body part trajectories, to identify images where the decoder might make large errors. We provide various frame-selection methods for this purpose. In particular the user can:

- select frames if the likelihood of a particular or all body parts lies below *p*_bound_ (note this could also be due to occlusions rather then errors).
- select frames where a particular body part or all body parts jumped more than *E* pixels from the last frame.
- select frames if the predicted body part location deviates from a state-space model [49, 50] fit to the time series of individual body parts. Specifically, this method fits an Auto Regressive Integrated Moving Average (ARIMA) model to the time series for each body part. Thereby each body part detection with a likelihood smaller than *p_bound_* is treated as missing data. An example fit for one body part can be found in Figure 6a. Putative outlier frames are then identified as time points, where the average body part estimates are at least *E* pixel away from the fits. The parameters of this method are *E, p*_bound_, the ARIMA parameters as well as the list of body parts to average over (can also be “all”).

All this can be done for a specific video by typing:

~~~
>> deeplabcut.extract_outlier_frames(config_path,[‘videofile_path’])
~~~

**FIG. 6.**
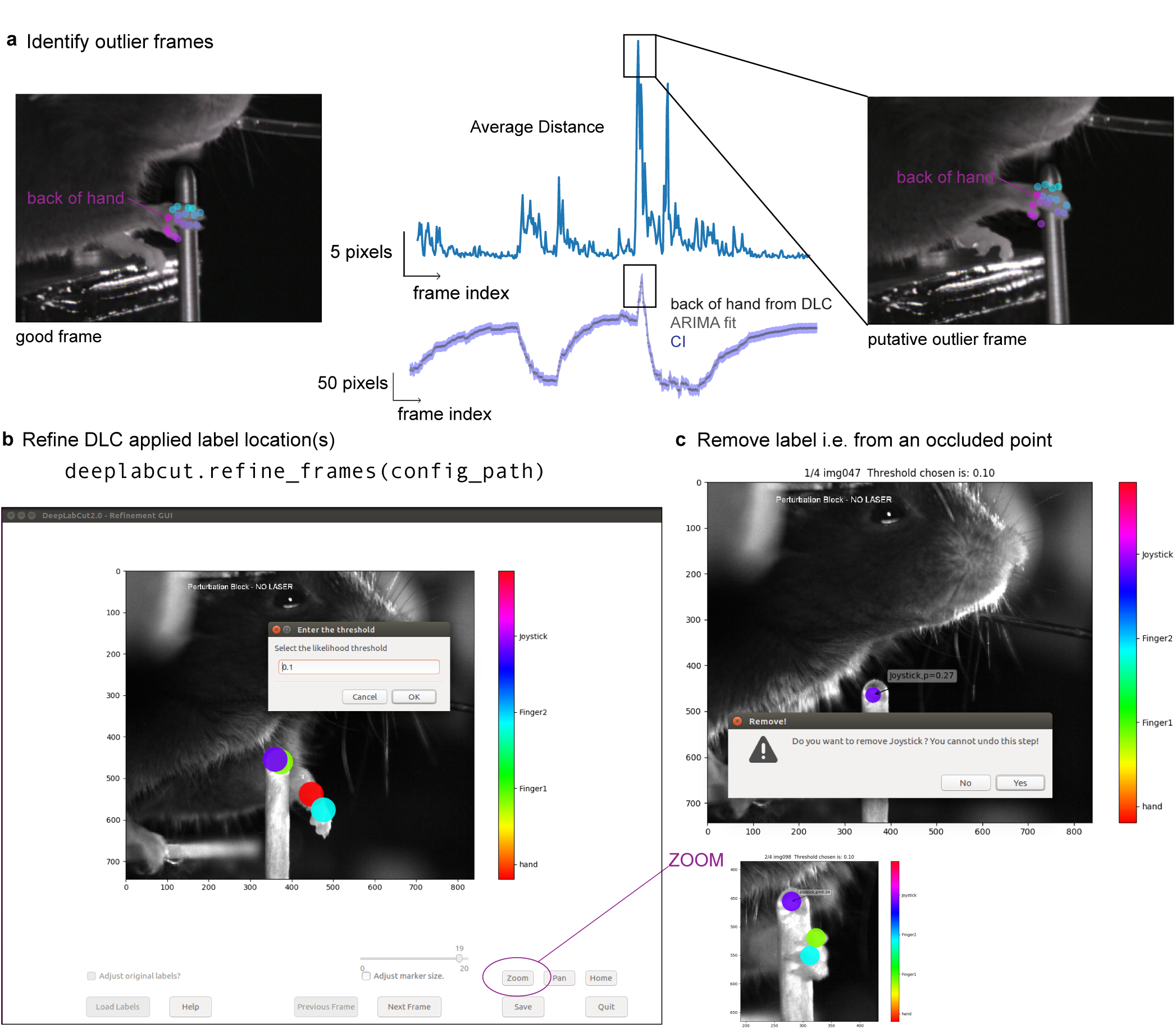
Refinement Tools. A user can refine the network by first extracting outliers and then by manually correcting the annotations based a dedicated graphical user interface (a) Outlier detection: For illustration we depict the x-coordinate estimate by DeepLabCut for the back of the hand and an ARIMA fit with 99%-confidence interval in blue. The average Euclidean distance for the tracked 17 body parts of the hand to the fit is also depicted. This distance can be used to find putative outliers as indicated by the corresponding frames. (b-c) Then adjust the labels and/or remove occluded labels.

In general, depending on the parameters, these methods might return much more frames than the user wants to extract (*numframes2pick*). Thus, this list is then used to select outlier frames either by randomly sampling from this list (‘uniform’) or by performing ‘k-means’ clustering on the corresponding frames (same methodology and parameters as in Step C). Furthermore, before this second selection happens, the user is informed about the amount of frames satisfying the criteria and asked if the selection should proceed. This step allows the user to perhaps change the parameters of the frame-selection heuristics first. The user can run the *extract_outlier_frames* iteratively, and (even) extract additional frames from the same video. Once enough outlier frames are extracted the refinement GUI can be used to adjust the labels based on user feedback (Step K). This function has many optional parameters, so a user should reference the help by typing deeplabcut.extract_outlier_frames? (and see Step N).

## K. Refine Labels: Augmentation of the Training Dataset

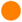**Timing**: The time required to refine the labels depends on the speed of the experimenter to identify the incorrect label and is mainly related to the number of body parts and the total number of extracted outlier frames.

Based on the performance of DeepLabCut, four scenarios are possible:

(A) Visible body part with accurate DeepLabCut prediction. These labels do not need any modifications.

(B) Visible body part but wrong DeepLabCut prediction. Move the label’s location to the actual position of the body part.

(C) Invisible, occluded body part. Remove the predicted label by DeepLabCut with a right click. Every predicted label is shown, even when DeepLabCut is uncertain. This is necessary, so that the user can potentially move the predicted label. However, to help the user to remove all invisible body parts the low-likelihood predictions are shown as open circles (rather than disks).

(D) Invalid images: In an unlikely event that there are any invalid images, the user should remove such an image and their corresponding predictions, if any. Here, the GUI will prompt the user to remove an image identified as invalid.

The labels for extracted putative outlier frames can be refined by opening the GUI:

~~~
>> deeplabcut.refine_labels(config_path)
~~~

This will launch a GUI where the user can refine the labels (Figure 6). The GUI is launched taking the size of the user’s screen into account, so if two monitors are used (in landscape mode) place ‘Screens=2’ after config_path (The default is 1). For other configurations, please see the linked troubleshooting guide (Section O). Use the ‘Load Labels’ button to select one of the subdirectories, where the extracted frames are stored. It is also possible to correct labels that have already been applied (or refined) by a user by simply check the ‘adjust original label?’ checkbox before loading the labels file. Every label will be identified by a unique color. For better chances to identify the low-confidence labels, specify the threshold of the likelihood. This changes the body parts with likelihood below this threshold to appear as circles and the ones above as solid disks while retaining the same color scheme. Next, to adjust the position of the label, hover the mouse over the labels to identify the specific body part, left click+drag it to a different location. To delete a specific label, right click on the label (once a label is deleted, it cannot be retrieved). After correcting the labels for all the frames in each of the subdirectories, the users should merge the data set to create a new dataset. The iteration parameter in the *config.yaml* file is automatically updated.

~~~
>> deeplabcut.merge_datasets(config_path)
~~~

Once the dataset is merged, the user can test if the merging process was successful by plotting all the labels (Step E). Next, with this expanded training set the user can now create a novel training set and train the network as described in Steps F and G. The training dataset will be stored in the same place as before but under a different ‘iteration #’ subdirectory, where the ‘#’ is the new value of ‘iteration’ variable stored in the project’s configuration file (this is automatically done).

If after training the network generalizes well to the data, proceed to Step I to analyze new videos. Otherwise, consider labeling more data (Section J).

## L. Anticipated Results

DeepLabCut has been used for pose estimation of various points of interest for different behaviors and animals (Fig 1). Due to the initial pre-training of the ResNet (i.e. the ResNet is first pre-trained on ImageNet [51]), as we have shown in our original study [12], DeepLabCut does not typically require many labeled frames to train on. We previously showed that even as little as 100 frames could be labeled for accurate tracking of a mouse snout in an open field-like scenario with challenging background and lighting statistics. The technical report also contains examples for hand articulation and fruit fly pose-estimation during egg-laying [12].

Here, we provide a new use case, where the experimenters wanted to quantitatively track the 3D kinematics of a cheetah performing a high-speed pursuit task as would be done during hunting in the wild [52]. This specific application has many challenges for any pose-estimation approach: highly variable scene statistics where the animal traverses over a field of grass and stirs up dust, different weather conditions, the size of the animal in the frame changes dramatically as it runs into a “camera trap” of 6 high-speed cameras (90 FPS, 1080p resolution), and the different cheetahs have diverse coats and highly articulated leg and body movements (Figure 5a,b). In total, we labeled 255 frames across multiple cameras (some of which come from the video in the figure) using the toolbox presented in this guide. Thus, the 3D kinematics can be reconstructed from one network being trained on multiple views, and the user only needs the camera calibration files to reconstruct their data. This camera calibration can be done in many programs, including OpenCV [36].

Specifically, we first selected 10 frames each from 12 different videos by using the uniform random sampling across the length of each video. Next, labeling was performed on the initial set using the supplied GUI. We trained with automatic cropping and rescaling; due to the large input size image of 1920 x 1080 pixels, we changed ‘max_input_size’ to 2000. After training, we iteratively refined images and re-trained the network until we had a set of 255 frames that allowed for reasonable performance and generalization. In Figure 7 we show how multiple cameras can be used to train a single network (Figure 7a), and an example image sequence with trajectories of past actions (Figure 7b), and using 2 cameras we show a 3D reconstruction [53, 54] of one body side (Figure 7c). This is an ongoing project studying the biomechanics of cheetah hunting and will be published elsewhere. Yet, we hope this example use case shows how DeepLabCut can be utilized even for very challenging, large video-resolution tracking problems “in the wild”, and empowers the user to apply this approach to their own unique and exciting experiments.

## M. upyter Notebooks for Demonstration of the DeepLabCut Work-flow

We also provide two Jupyter notebooks for using DeepLabCut on both a pre-labeled dataset, and on the end user’s own dataset. Firstly, we prepared an interactive Jupyter notebook called *run_yourowndata.ipynb* that can serve as a template for the user to develop a project. Furthermore, we provide a notebook for an already started project with labeled data. The example project, named as *Reaching-Mackenzie-2018-08-30* consists of a project configuration file with default parameters and 20 images, which are cropped around the region of interest as an example dataset. These images are extracted from a video, which was recorded in a study of skilled motor control in mice [55]. Some example labels for these images are also provided. The notebooks can be started by typing in the terminal:

~~~
jupyter-notebook Demo-labeledexample-MouseReaching.ipynb
jupyter-notebook Demo-yourowndata.ipynb
~~~

## N. Overview of Commands

Here is a ‘quick-guide’ of the minimal commands used for running DeepLabCut in Python/ipython. All these functions have additional optional parameters that the user may change. The user can invoke ‘help’ for any function and get more information about these optional parameters as follows:

~~~
In ipython/Jupyter Notebooks: deeplabcut.nameofthefunction?
In Python: help(deeplabcut.nameofthefunction)
~~~

**TABLE I:** Summary of Commands

| Steps | Command |
| --- | --- |
| Open ipython or Python<br>import deeplabcut | ipython (or python)<br>import deeplabcut |
| Create a new project.<br>See section A for details | **Python/ipython:**<br>>> deeplabcut.create_new_project(‘project name’, ‘experimenter’,<br>[‘path of video 1’, ‘path of video2’, ...]) |
| Set a config_path<br>for ease of use | >> config_path = ‘yourdirectory/your_deeplabcut/config.yaml’ |
| Extract frames<br>See section C for details | **Python/ipython:**<br>>> deeplabcut.extract_frames(config_path, mode= ‘automatic’) |
| Label frames<br>See section D for details | **Python/ipython:**<br>>> deeplabcut.label_frames(config_path) |
| Check labels<br>[OPTIONAL]<br>See section E for details | **Python/ipython:**<br>>> deeplabcut.check_labels(config_path) |
| Create training dataset<br>See section F for details | **Python/ipython:**<br>>> deeplabcut.create_training_dataset(config_path) |
| Train the network<br>See section G for details | **Python/ipython:**<br>`>> deeplabcut.train_network(config_path)` |
| Evaluate the trained network<br>See section H for details | **Python/ipython:**<br>`>> deeplabcut.evaluate_network(config_path)` |
| Video analysis and plotting Results<br>See section I for details | **Python/ipython:**<br>`>> deeplabcut.analyze_videos(config_path,`<br>`[‘path of video 1’, ‘path of video2’, ...])` |
| Video analysis and plotting Results<br>See section I for details | **Python/ipython:**<br>`>> deeplabcut.create_labeled_video(config_path,`<br>`[‘path of video 1’, ‘path of video2’, ...])` |
| Video analysis and plotting Results<br>See section I for details | **Python/ipython:**<br>`>> deeplabcut.plot_trajectories(config_path,`<br>`[‘path of video 1’, ‘path of video2’, ...])` |
| Refinement: Extract outlier frames<br>See section J for details | **Python/ipython:**<br>`>> deeplabcut.extract_outlier_frames(config_path, [‘path of video 1’,`<br>`‘path of video 2’])` |
| Refine Labels: Augmentation of the training dataset<br>See section K for details | **Python/ipython:**<br>`>> deeplabcut.refine_labels(config_path)` |
| Combine datasets: Augmentation of the training dataset<br>See section K for details | **Python/ipython:**<br>`>> deeplabcut.merge_datasets(config_path)` |

## O. Troubleshooting

The Jupyter Notebooks provided on GitHub have been already executed so the user can see the expected outputs. They are also annotated with potential errors and considerations. Additionally, when running the Notebooks on Colaboratory (https://colab.research.google.com) the user can directly search for any error messages that arise, as they can involve TensorFlow or other packages not managed by us. In the DeepLabCut code we strived to make error messages intuitive for each step, and each output directs the user to the next step in the process.

There is also an actively maintained GitHub repository that has an “Issues” section for users to look for solutions to problems others encountered or report new ones, as well as a wiki page with FAQ (https://github.com/AlexEMG/DeepLabCut/wiki/Troubleshooting-Tips).

## Acknowledgements

DeepLabCut is an open-source tool on GitHub and has benefited from suggestions and edits by many individuals including Ronny Eichler, Jonas Rauber, Richard Warren, Taiga Abe, Hao Wu, and Jonny Saunders. In particular, the authors thank Ronny Eichler for input on the modularized version. The authors thank members of the Bethge Lab for providing the initial version of the Docker container. We also thank Meng Li, Jennifer Li & Drew Robson for use of the zebrafish image, Byron Rogers for use of the horse images, and Kevin Cury for the fly images. The authors are grateful to Eldar Insafutdinov and Christoph Lassner for suggestions on how to best use the TensorFlow implementation of DeeperCut. We also thank Adrian Hoffmann, Pranav Mamidanna and Gary Kane for comments throughout this project. Lastly, the authors thank the Ann van Dyk Cheetah Centre (Pretoria, South Africa) for kindly providing access to their cheetahs.

## Contributions

Conceptualization: AM, TN, MWM. AM, TN and MWM wrote the code. AP provided the cheetah data; AC labeled the cheetah data; AC, AM and AP analyzed the cheetah data. AM, MWM and TN wrote the manuscript with inputs from all authors. MWM and MB supervised the project.

